# Expression of the locus of enterocyte effacement genes during the invasion process of the atypical enteropathogenic *Escherichia coli* 1711-4 strain of serotype O51:H40

**DOI:** 10.1101/2024.02.01.578415

**Authors:** Fabiano T. Romão, Ana C. M. Santos, Vanessa Sperandio, Rodrigo T. Hernandes, Tânia A. T. Gomes

## Abstract

Atypical enteropathogenic *Escherichia coli* (aEPEC) is a significant cause of diarrhea in developing countries. Some aEPEC strains, including the Brazilian representative strain of serotype O51:H40 called aEPEC 1711-4, can use flagella to attach to, invade, and persist in T84 and Caco-2 intestinal cells. They can even translocate from the gut to extraintestinal sites in a rat model. Although various aspects of the virulence of this strain were studied and the requirement of the T3SS for the efficiency of the invasion process was demonstrated, the expression of the LEE genes during the invasion and intracellular persistence remains unclear. To address this, the expression of flagella and the different LEE operons was evaluated during kinetic experiments of the interaction of aEPEC 1711-4 with enterocytes *in vitro*. The genome of the strain was also sequenced. The results showed that flagella expression remained unchanged, but the expression of *eae* and *escJ* increased during the early interaction and invasion of aEPEC 1711-4 into Caco-2 cells, and there was no change 24 hours post-infection during the persistence period. The number of pedestal-like structures formed on HeLa cells also increased during the 24-hour analysis. No known gene related to the invasion process was identified in the genome of aEPEC 1711-4, which was shown to belong to the global EPEC lineage 10. These findings suggest that LEE components and the intimate adherence promoted by intimin are necessary for the invasion and persistence of aEPEC 1711-4, but the detailed mechanism needs further study.

**Importance:** Atypical enteropathogenic *Escherichia coli* (aEPEC) is a major cause of diarrhea, especially in developing countries, like Brazil. However, due to the genome heterogeneity of each clonal group, it is difficult to comprehend the pathogenicity of this strain fully. Among the aEPEC strains, 1711-4 can invade eukaryotic cells *in vitro*, cross the gut barrier, and reach extraintestinal sites in animal models. By studying how different known aEPEC virulence factors are expressed during the invasion process, we can gain insight into the commonality of this phenotype among other aEPEC strains. This will help us develop preventive measures to control infections caused by invasive strains. No known virulence-encoding genes linked to the invasion process were found. Nevertheless, additional studies are still necessary to evaluate the role of other factors in this phenotype.

## Introduction

Enteropathogenic *Escherichia coli* (EPEC) is still an important pathogen related to diarrheal diseases in developing countries. In Brazil, it is the most frequent *E. coli* pathotype isolated from diarrhea (1). The EPEC pathotype is subdivided into two subgroups, typical and atypical, based on the production of the adhesion factor named bundle forming pilus (BFP), present on typical EPEC (tEPEC) and absent on atypical EPEC (aEPEC) (2). Due to the absence of BFP, aEPEC requires a prolonged time of interaction with enterocytes *in vitro* to adhere to and promote attaching and effacing (A/E) lesion (2) that is mediated by proteins encoded by genes present in the Locus of Enterocyte Effacement (LEE) pathogenicity island. The A/E lesion is characterized by the intimate adherence between bacteria and host cells, which promotes microvilli reshuffling into pedestal-like structures. Such cell surface modification is promoted by the injection of diverse proteins encoded in the LEE region through the type three secretion system (T3SS), which result in the mobilization of actin and other cytoskeletal proteins to the adherence site (2–4).

The LEE comprises five polycistronic operons (LEE1, LEE2, LEE3, LEE4, and LEE5), two bicistronic operons (*espG-rorf1* and *grlA-grlR*), and four independent genes (*etgA*, *cesF*, *map*, and *escD*), and together with diverse LEE-independent genes, are directly involved in the A/E lesion formation and diarrhea caused by the EPEC pathotype (5, 6).

The aEPEC subgroup is genetically heterogeneous (4, 7–9) and different strains may carry additional virulence factors that might contribute to the initial stages of enterocyte colonization and diarrhea. Strains belonging to the O51:H40 serotype comprise one of the most frequent among aEPEC isolated from diarrhea in Brazil (1, 10–13). One such strain, aEPEC 1711-4 was used in various previous studies focused on enlarging our knowledge of the virulence of the aEPEC strains. In these studies, we showed that this strain could use flagella to attach to enterocytes during the initial colonization (14), and was able to invade and persist inside T84 and Caco-2 cells (15), induce IL-8 (14, 15) and translocate from the gut to different extraintestinal sites causing systemic infection in a rat model (16). Sampaio et al. (17) showed that the 1711-4 isogenic strain lacking the T3SS was not able to invade Caco-2 cells properly, to translocate from the gut to extraintestinal sites or promote A/E lesion in a rabbit ileal loop model, demonstrating the relevance of this system in all these phenotypes. Although various aspects of the virulence of this strain were studied and the requirement of the T3SS for the efficiency of the invasion process was demonstrated, the impact of the LEE genes expression during the invasion and intracellular persistence remains unclear; additionally, full details regarding the 1711-4 strain’s genome are unknown. Therefore, in this study, we evaluated the expression of the LEE regions and flagella during the invasion and persistence process of aEPEC 1711-4 into Caco-2 and HeLa cells. We also provided the whole sequence and analyses of the aEPEC 1711-4 genome.

## Methods

### Bacterial strain

The aEPEC 1711-4 strain (serotype O51:H40) was isolated from a child with diarrhea during an epidemiological study on diarrhea, which was conducted in 1989 at the Universidade Federal de São Paulo (UNIFESP), Brazil (13). One fluorescent variant of aEPEC 1711-4 was obtained by transforming the wild-type strain with the recombinant plasmid pDP151 (Invitrogen), which encodes the fluorescent protein mCherry and confers resistance to ampicillin.

### Cell culture

Caco-2 cells ATCC® HTB-37™ were used to evaluate differential expression of the LEE genes during the invasion process and intracellular persistence. The cells were cultivated in 24-well plates using Dulbecco’s Modified Eagle Medium (DMEM, Gibco, USA) supplemented with 10% Fetal bovine serum (Gibco, USA), 1% antibiotic mixture (penicillin-10,000 U/ml and streptomycin-10 mg/ml, ThermoFisher, USA), and 1% non-essential amino acids mixture (Life Technologies, USA), in an atmosphere of 5% CO_2_ at 37°C for up to 10 days, to enable cell polarization and differentiation. Before the assays, the monolayer was washed three times with phosphate-buffered saline (PBS), and fresh DMEM supplemented with 2% FBS was added.

HeLa cells stably expressing Lifeact::GFP (18, 19) were used to evaluate the formation of pedestal-like structures and invasion in HeLa cells. The cells were grown in DMEM with 10% FBS, 1× PSN, 50 µg/ml of hygromycin B (ThermoFisher, USA), and kept in a 5% CO_2_ atmosphere at 37 °C. For the fluorescence microscopy assay, HeLa(actin-GFP) cells (5×10^5^ cells/per dish) were seeded onto 35 mm cell culture dishes with glass bottom (Corning, USA) 48-h before the assay.

### Evaluation of the bacterial invasion and intracellular persistence

The invasion assay was performed as described by Pacheco *et al*. (20) with modifications. Briefly, overnight cultures grown in Lysogeny broth (LB) were adjusted to ∼ 0.5 OD_600_, inoculated in a ratio of 1:50 in two 24-well plates containing polarized and differentiated Caco-2 cells, and incubated at 37 °C for 1.5 h, 3 h, and 6 h. After the incubation period, one plate was washed three times with PBS, 1 ml of DMEM containing 2% FBS. Then, 100 µg/ml of gentamicin was added to each well and plates were incubated for 1 hour to kill extracellular bacteria. The other plate was washed and kept untreated. After incubation, cells in both plates were lysed with 1% Triton X-100, serially diluted, plated onto MacConkey agar plates, and incubated at 37°C for 18 h to determine the numbers of total bacteria (TB) and intracellular bacteria (IB). The invasion index was calculated as (IB × 100 / TB). The assays were performed in biological and technical triplicates, and results were presented as mean ±standard deviation.

The evaluation of bacterial persistence was performed as the invasion assay. However, after 6 h of incubation, the cells were washed three times with PBS, 1 ml of DMEM supplemented with 2% FBS and 100 µg/ml of gentamicin was added to each well, and the preparations were incubated at 37°C for 18 h, totalizing a 24 h assay. After incubation, the monolayers were washed with PBS, lysed with Triton X-100, serially diluted, and plated onto MacConkey.

### Fluorescence actin staining (FAS) assay and F-actin pedestal quantification

The assay was performed as previously published. Briefly, aEPEC 1711-4 expressing mCherry was grown statically in LB for 18 h at 37°C. HeLa (Actin-GFP) cells in DMEM supplemented with 2% FBS and 100 µg/ml of ampicillin were infected with 1.5×10^7^ bacteria prepared from an overnight culture. Pedestal formation was evaluated after 1.5 h, 3 h, and 6 h of incubation at 37 °C in an atmosphere of 5% CO_2_. The assay was washed with PBS and fixed with 3.7% formaldehyde. Then, the coverslips were washed with PBS, incubated with 8 µM of FITC-phalloidin (Invitrogen), washed with PBS and Saline-Sodium Citrate buffer (SSC) [2×], treated with 100 µg/ml RNAseA (Sigma-Aldrich), washed with SSC 2×, incubated with 1.7 µM of propidium iodide, and rewashed with SSC 2×. The cells were visualized with a Zeiss confocal microscope with a 63×1.40 N.A. immersion oil objective. Pedestals were quantified by randomly imaging different fields while recording the number of cells showing F-actin accumulation foci. Results were presented as means of percentage (%) of infected cells with F-actin accumulation or number of pedestals per cell ±standard deviation. The assay was assessed in replicates and at least two independent experiments.

### qRT-PCR

The expression of the *ler, escJ, escV, escN, eae,* and *espA* genes, representing the different operons in the LEE region, and *fliC*, encoding the bacterial flagellin, was evaluated during the infection and persistence assays by qRT-PCR. For that, invasion and persistence were performed using Caco-2 cells seeded into 6-well plates. The monolayer was lysed at each specific time for invasion (90 min, 3 h, and 6 h) and persistence (6 h and 24 h), and total RNA was extracted using the RNeasy Plus Mini kit (Ambion, Lithuania). The RNA was quantified using a Biophotometer (Eppendorf, Germany), DNA was removed using DNAse (Invitrogen, USA), and the reverse transcription was performed using Superscript FirstStrand Synthesis for RT-PCR (Invitrogen, CA, USA). cDNA was amplified using the Master Mix kit (Thermofisher, USA) with specific primers designed (Table 1). *rpoA* was used as a gene expression control.

**Table 1:**
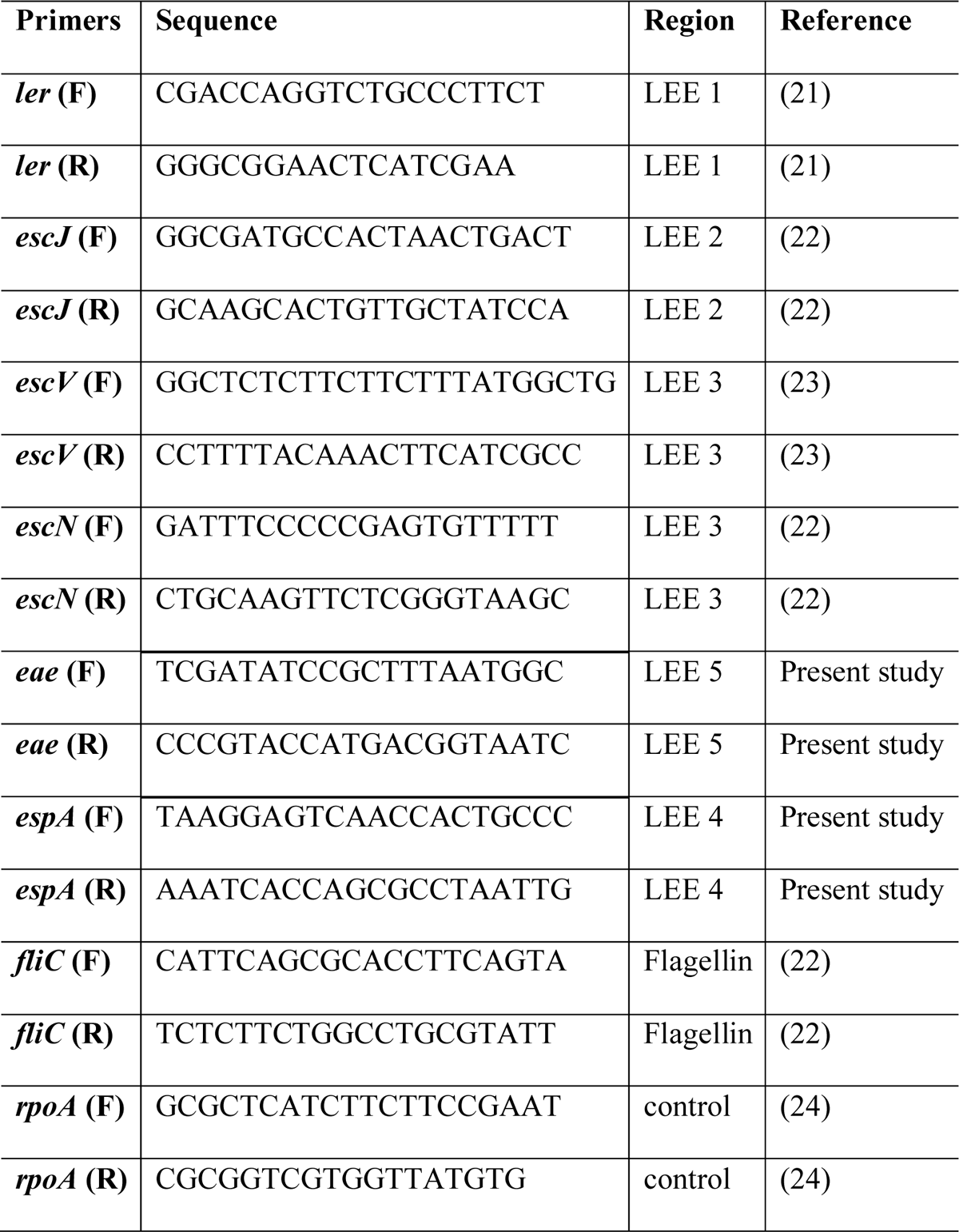
Primers used in RT-PCR.

### Whole genome sequencing and genomic analyses

The whole genome sequence of 1711-4 was performed using the PacBio RS sequencing System (Pacific Bio). The reads were assembled *de novo* using Canu assembler v 1.7.1 (25) and polished using Racon v1.4.13 (26). The assembled genome was used to perform diverse analyses at the Center for Genomic Epidemiology (CGE: http://genomicepidemiology.org/) using services for the identification of virulence genes (VirulenceFinder version 2.0)(27, 28), serotype (SeroTypeFinder version 2.0)(29), antibiotic resistance genes (ResFinder version 4.1)(30, 31), plasmids (PlasmidFinder version 2.0)(32), and sequence type determination (MLST version 2.0)(33), following the Warwick scheme. The virulence factors were also searched using the virulence factors database (VFDB)(34).

The Similar genome finder service with a threshold of 0.001 and distance of 0.01 was used to identify the published genomes like 1711-4 to identify to which Clonal complex the strain belonged. The MLST of the 50 *E. coli* strains similar to 1711-4 were evaluated in the CGE as described above. The clonal complex of the strains was searched using EnteroBase (https://enterobase.warwick.ac.uk/) (35). To understand their relationship, a phylogenetic tree was built using the Bacterial Genome Tree Service (36) with all the strains. To build the tree, prototype *E. coli* O127:H6 str. E2348/69 (ST15-B2), O157:H7 str. Sakai (ST11-E), O104:H4 str.2011C-3493 (ST678-B1), and H10407 (ST10-A) were used as outgroups.

Another phylogenetic tree was built to identify the clonal relationship of 1711-4 with EPEC and EHEC clonal lineages previously published (9, 37) at the BV-BCR. The treeś final layouts were built using iTOL v 6.8 (38).

### Statistical analyses

The results were compared and evaluated using the non-parametrical t-student test. The numbers of FAS-positive signals and pedestals were compared using One-way ANOVA followed by the post hoc Turkey HSD test. *P* values ≤0.05 were considered statistically significant. The statistical analyses were performed using Prism GraphPad ver. 8.4.2.

## Results

### Adhesion and invasion of aEPEC 1711-4 with epithelial cells occur at different time points

A kinetic interaction was performed to understand the 1711-4 adhesion and invasion behaviors in the colonization of Caco-2 cells. The number of bacteria interacting with Caco-2 cells increased from 1.5 h to 3 h (*P* ≤ 0.0001) but did not change between 3 h and 6 h (Figure 1A).

**Figure 1.**
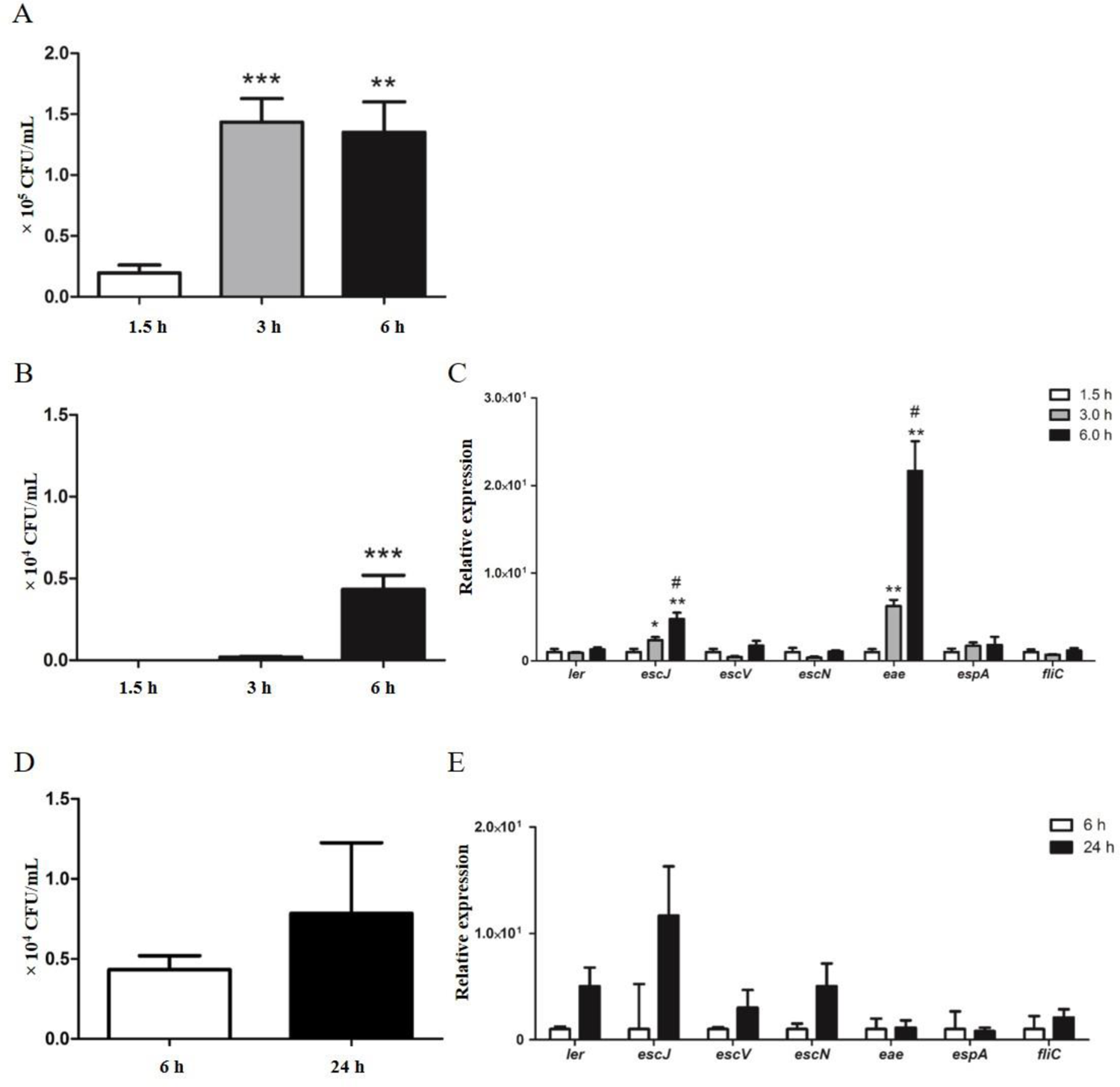
Kinetic analyses of the interaction, invasion, and LEE gene and *fliC* expression during aEPEC 1711-4 incubation with polarized and differentiated Caco-2 cells. (A) kinetics of total interaction, showing that the numbers of bacteria interacting with Caco-2 cells in 3 and 6 hours were significantly higher than in 1.5 h, while no differences were observed between 3 and 6 h; (B) kinetics of invasion, showing that invasiveness could be observed only after 3 h, being significant after 6 h of interaction; (C) Relative expression of genes representing different LEE regions and flagella during the initial adherence and invasion processes; the expression of the *escJ* and *eae* genes increased from 1.5 h to 3 h and from 3 h to 6 h; (D) kinetics of the bacterial intracellular persistence; no significant difference in bacterial counting was identified between 6 h and 24 h; (E) Relative gene expression of genes during the persistence; although expression of the *ler, escJ,* and *escN* genes increased, no significant difference was identified between 6 h and 24 h. * *P* ≤ 0.05, ** *P ≤* 0.01 and *** *P* ≤ 0.0001; for relative gene expression # *P ≤* 0.05 comparing 3 h and 6 h.

Although the total interaction did not change, the quantification of the invasion efficiency in the same period showed that the aEPEC 1711-4 invasion process occurred only after 1.5 h and increased after 3 h of interaction (Figure 1B).

### Expression of *eae* and *escJ* increases during the invasion process in Caco-2 cells but does not change during the intracellular persistence

As the transcription levels of the LEE genes during the initial interaction of aEPEC 1711-4 with Caco-2 cells is unknown, the *ler, escJ, escV, escN, eae,* and *espA* transcription levels were evaluated to verify the impact of their operons in the process. Additionally, as it is known that flagella participate in the initial interaction of 1711-4 with Caco-2 cells (14), the transcription of *fliC*, encoding the flagellin, was also evaluated.

The transcription of the *escJ* and *eae* genes from aEPEC 1711-4 gradually increased during the first 6 hours of interaction with Caco-2 cells, while the transcription of the other genes evaluated did not change along the period (Figure 1C).

During the bacterial intracellular persistence, although *ler*, *escJ*, and *escN* displayed higher levels of transcription in 24 h than in 6 h, there was no notable change in expression of the genes, comparing 6 h and 24 h (Figure 1E). Similarly, the number of bacteria inside the Caco-2 cells barely changed from 6 h to 24 h (Figure 1D).

### Pedestal formation and bacterial invasiveness increase along the interaction with HeLa cells

The number of pedestals per cell increased gradually from 3 h to 6 h (Figure 2, Figure 3A), being significantly higher at 6 h (Figure 3A). In the same way, the invasion of HeLa cells reached 12% at 6 h but internalized bacteria were scarce 3 h post-infection (Figure 3B).

**Figure 2.**
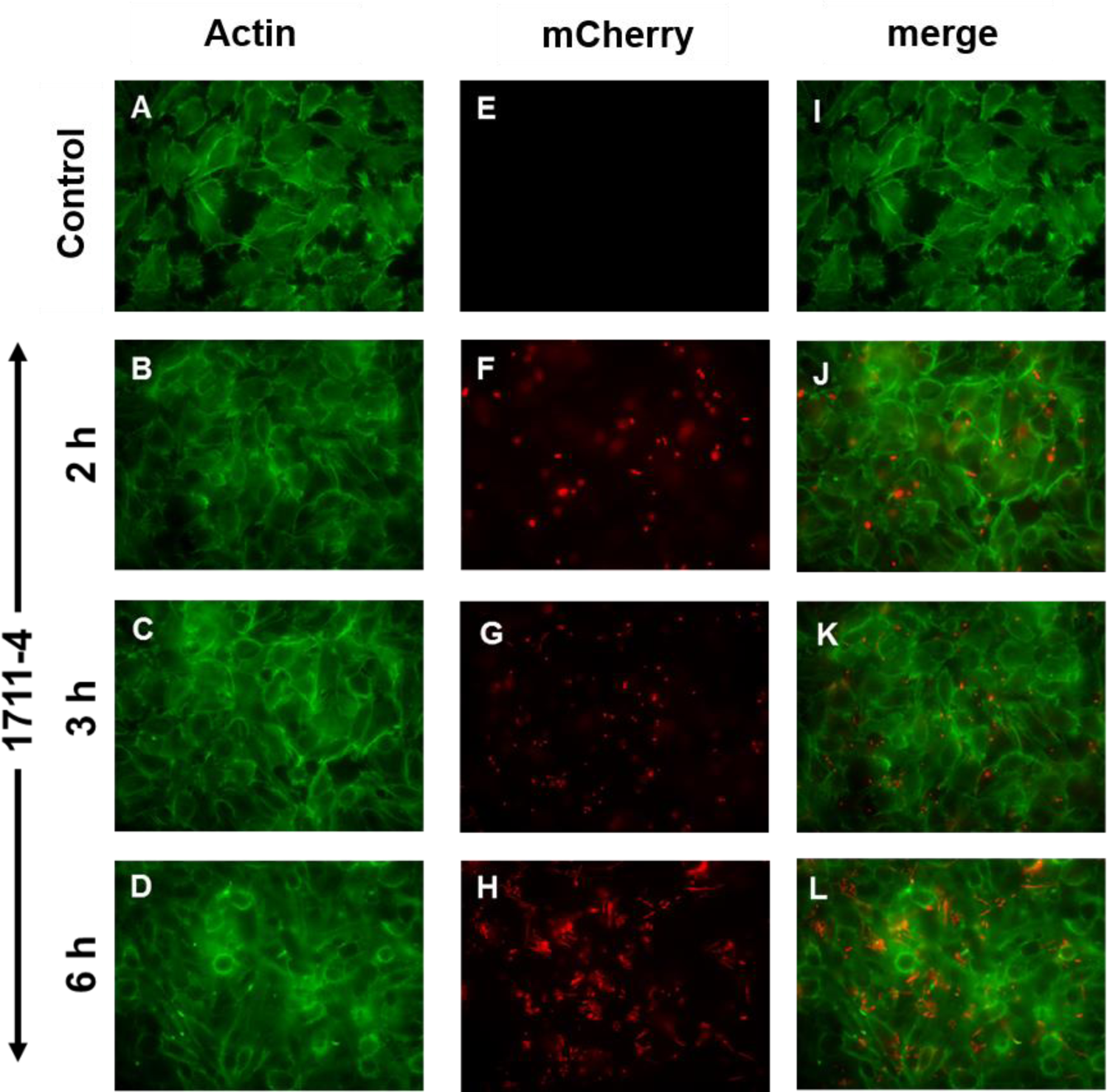
Kinetic of pedestal formation of aEPEC 1711-4 in HeLa cells. The pedestal formation was evaluated from 2 to 6 h of interaction. From 3 h to 6 h, the number of bacteria interacting with HeLa cells and the number of pedestal structures per cell increased.

**Figure 3.**
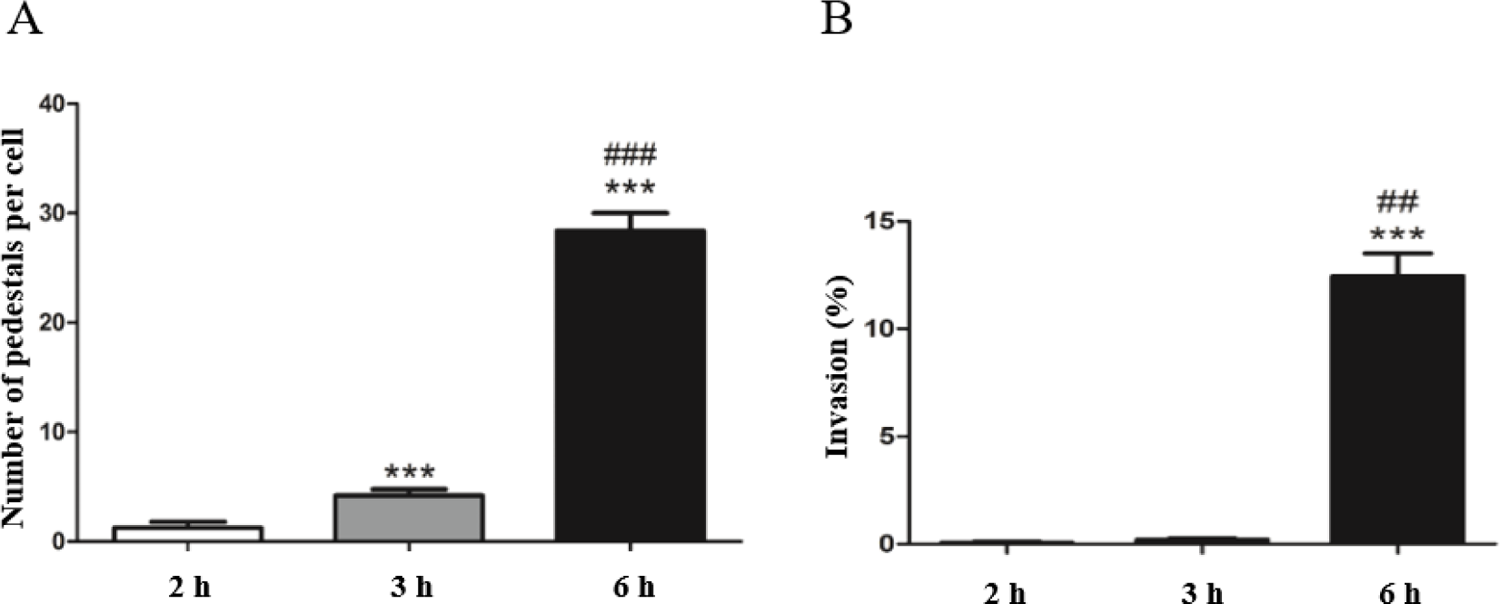
Quantification of pedestal formation and invasion rates in HeLa cells. (A) The number of pedestals per cell significantly increased from 2 h to 3 h and from 3 h to 6 h post-infection. (B) The invasion rate was significantly high only after 6 hours post-infection. (*) was used to express differences between 2 h against 3 h or 6 h, while (#) was used for comparison between 3 h and 6 h. * *P* ≤ 0.05, ** *P ≤* 0.01 *** *P* ≤ 0.0001, ## *P ≤* 0.01, and ### *P* ≤ 0.0001.

### aEPEC 1711-1 belongs to the EPEC global lineage 10, ST10 complex, and phylogroup A

The whole genome of aEPEC 1711-4 comprised 4,722,189 bp and belonged to phylogroup A. Also, the serotype O51:H40 was confirmed *in silico* but the MLST ST was not identified.

To detect to which clonal complex the strain belonged, Mash/MinHash was used to screen the genomes similar to aEPEC 1711-4 and deposited at the NCBI. The strains identified belonged mainly to the ST10 (43/51), two belonged to the clonal complex of sequence type 10 (ST10 cplx) (ST34 and ST752), and six strains belonged to underrepresented STs (ST5332, ST5339, and ST5353) not assigned to any clonal complex but that clustered together with the strain from ST10 cplx. Most strains identified in the cluster belonged to the aEPEC pathotype and were isolated worldwide (Figure 4 and Supp Table 1).

**Figure 4.**
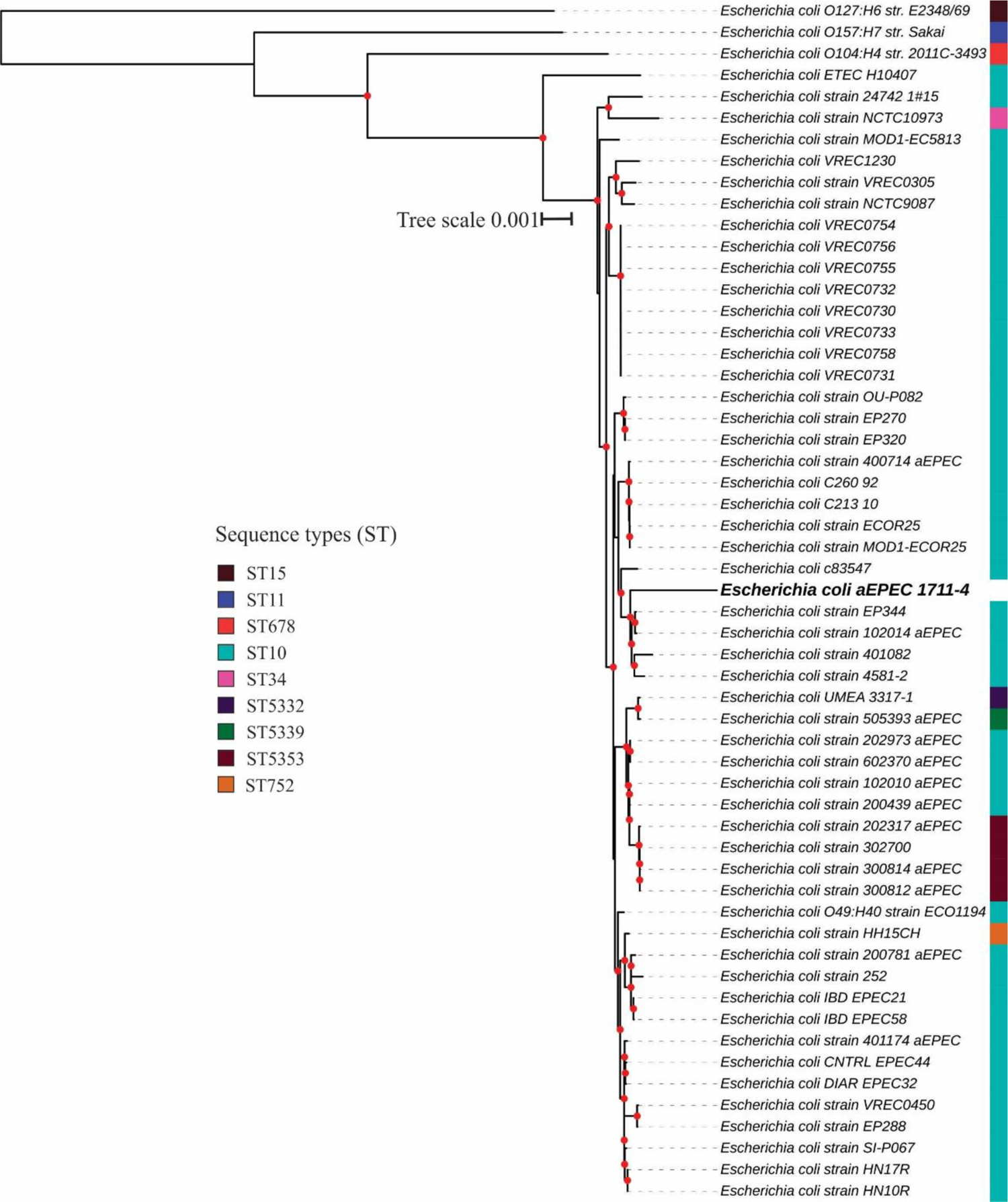
aEPEC 1711-4 similar genomes codon tree. Red dots in the nodes represent bootstrap ≥90. The *E. coli* prototype strains O127:H6 str. E2348/69 (ST15-B2), O157:H7 str. Sakai (ST11-E), and O104:H4 str. 2011C-3493 (ST678-B1), and ETEC str. H10407 (ST10-A) were used as outgroups.

Additionally, the evaluation of the EPEC/EHEC global clonal lineages showed that aEPEC 1711-4 belongs to the EPEC10 clonal group (Figure 5). Regarding virulence factors, aEPEC 1711-4 carried a complete LEE region and the intimin subtype theta. The non-LEE effectors NleB1, NleB2, NleE1, and NleH1 and the adhesins ECP, ELF, FdeC, YDF, and HCP were also identified in the genome. No known invasin, plasmid, or genes related to extraintestinal pathogenicity or antimicrobial resistance were identified in the aEPEC 1711-4 genome.

**Figure 5.**
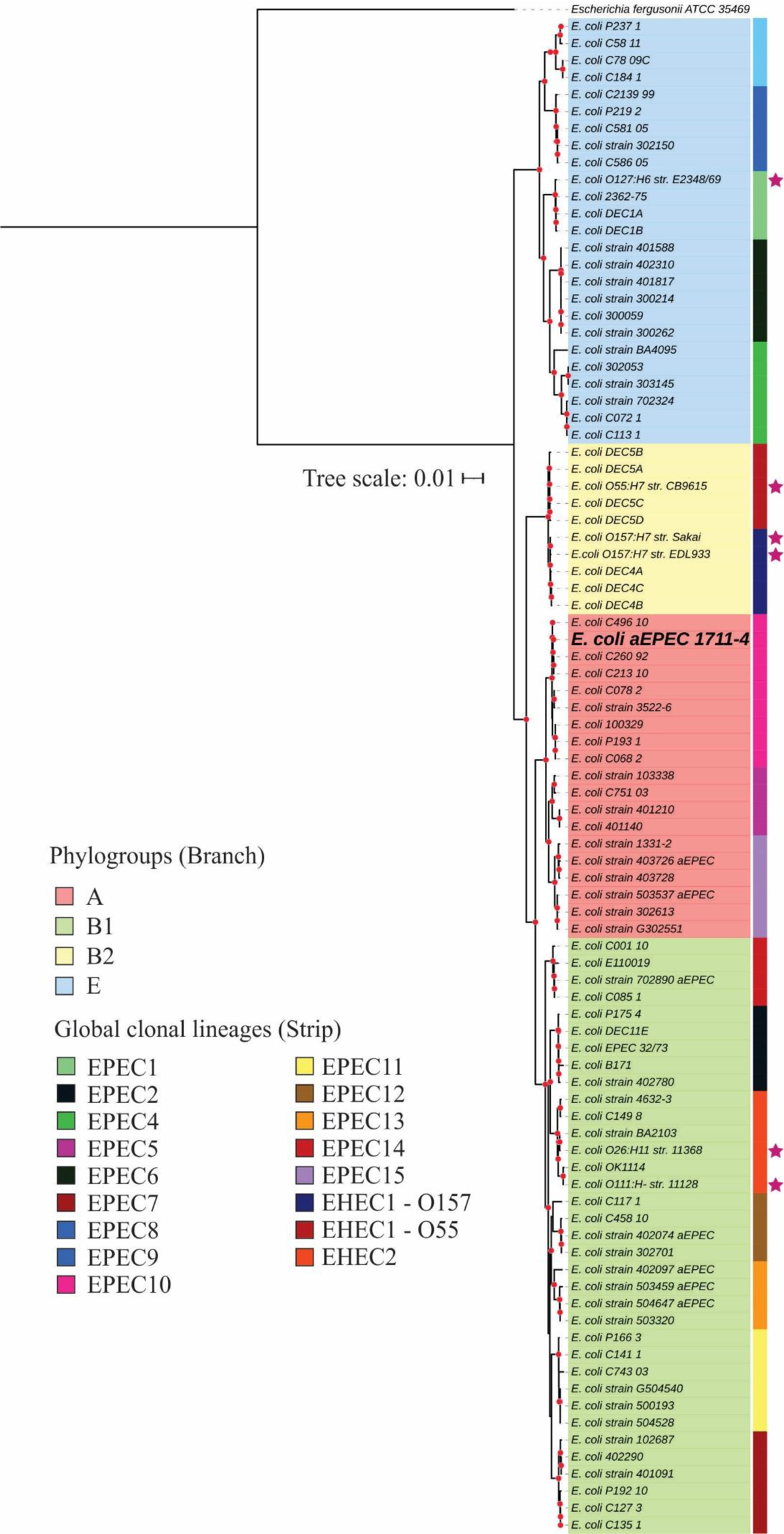
Relationship of aEPEC 1711-4 and the EPEC and EHEC global lineages. *Escherichia fergusonii* ATCC35469 was used as an outgroup to root the tree. A pink star in the tree indicates EPEC and EHEC prototype or representative strains.

## Discussion

The aEPEC pathotype comprises genetically heterogeneous bacteria that harbor diverse accessory genes that contribute to their virulence (7–9). In contrast with the tEPEC pathotype, the mechanisms related to aEPEC pathogenicity are not well established due to their considerable diversity. Furthermore, aEPEC is still one of the major causes of diarrhea in developing countries like Brazil, being the most prevalent diarrheagenic pathotype isolated from diarrheal diseases (1).

In the present work, we enlarged the knowledge regarding the 1711-4 strain, one representative Brazilian aEPEC strain belonging to the O51:H40 serotype. As the expression of flagella and the LEE PAI genes in the invasion and intracellular bacterial persistence was unknown, we evaluated the expression of five LEE genes, representing the different LEE regions, and flagella during the early invasion and persistence processes in Caco-2 cells.

During the interaction of aEPEC 1711-4 with Caco-2 cells, the number of associated bacteria reached its maximum at 3 h, while invasiveness was identified only after 3 h. During this process, the *escJ* and *eae* genes continuously increased their expression rate, reaching their maximum 6 h post-infection. This increase suggested that the proteins encoded by these genes might be involved in the invasion process. The *eae* gene codes for the adhesin intimin, responsible for the bacterial intimate adherence to host cells and represents the LEE 5 region.

In contrast, *escJ* represents the LEE 2 region and codes for one of the internal membrane structural proteins of the type 3 secretion system (T3SS), which is responsible for the injection of all effectors involved in A/E lesion formation, including the secretion of Tir, the intimin receptor that is also encoded in the LEE 5 region (2, 21, 39).

The T3SS is an essential virulence factor related to the pathogenicity of diverse bacterial genera like *Shigella*, *Salmonella,* and *Yersinia* (39–42). Studies showed that it is expressed constitutively in some pathogens like *Shigella*, although the T3SS proteins are only produced when the bacteria interact with the host cell membrane (43–45). Additionally, the T3SS expression is repressed once *Shigella* is released in the cytosol (43).

During the interaction with HEp-2 cells, it was observed that the expression of LEE operons in the tEPEC prototype strain E2348/69 was different from that of aEPEC 1711-4 infection of Caco-2 cells (46). In tEPEC, the *ler* gene was repressed at 3 hours post-infection, while the expression of *tir* and *eae* increased after 3 hours, and *espA* after 5 hours. However, there was no notable change in the expression of *escV*. In contrast, during aEPEC 1711-4 infection of Caco-2 cells, no significant change in the expression of the *ler*, *escN*, and *espA* genes was found during the studied period of up to 24 hours.

Some results were similar to those reported with tEPEC (46), such as the absence of significant change in *escV* expression and an increase in *eae* expression at 3 hours post-infection. However, in tEPEC, no increase in *eae* expression was identified after 5 hours, while in aEPEC 1711-4, *eae* expression kept increasing up to 6 hours post-infection and maintained the expression rate until 24 hours. It is interesting to note that in *E. albertii,* another invasive pathogen that harbors the PAI LEE, the expression of the LEE regions also differed from that identified in the present study. In the *E. albertii* strain 1551-2, there was no difference in the expression of any of the evaluated genes (*ler, escJ, escV, escN, eae,* and *espA*) at 1.5 h, 3 h, and 6 h post-infection (22), while all genes but *espA* showed an increase on gene expression at 24 h post-infection (22). This comparative and complex scenario perhaps indicates that the expression of the LEE genes can be differentially regulated in different isolates, and perhaps this fact can lead to differences in their virulence potential.

In aEPEC 1711-4, the differences in the gene expression identified may be linked to the invasion and/or persistence processes. In a previous study, it was demonstrated that the aEPEC 1711-4 isogenic strain lacking T3SS is unable to invade eukaryotic cells. Now, we have found that in the wild type 1711-4 strain, the *escJ* and *eae* genes are upregulated during the invasion process, and this expression remains high during the persistence stage. This supports the notion that LEE components play a role in the invasion process. Further analysis is needed to determine the role of individual LEE components in the invasion process of aEPEC 1711-4. Additionally, we confirmed that different from other LEE harboring invasive strains, aEPEC 1711-4 cannot multiply intracellularly but remained viable up to 48 h post-infection without an increase in the intracellular population (14, 20).

Our group previously showed that the flagella of aEPEC 1711-4 contribute to the early steps of bacterial adherence (14). For EHEC strains, it has been demonstrated that the *fliC* gene is repressed during the infection process (47). In the present study, we showed that *fliC* expression is continuous up to 24 h after infection, indicating that *fliC* continues to be expressed even after bacterial adherence to the cell surface for unknown reasons.

It was recently discovered that the 1711-4 strain lacks plasmids and belongs to the ST10 complex and EPEC 10 global lineage. This lineage is associated with EPEC strains that cause diarrhea and extraintestinal infections. The global EPEC 10 lineage, to which 1711-4 belongs, represents about 15% of all aEPEC strains isolated worldwide and is the most significant global lineage in phylogroup A. The genome of the 1711-4 strain was evaluated, and it was found to contain non-LEE effectors (NleB1, NleB2, NleE1, and NleH1) as well as five adhesins (ECP, ELF, FdeC, YDF, and HCP). However, no invasin or genes related to extraintestinal pathogenicity were identified that could explain its ability to invade cells and move from the gut to extraintestinal sites. The role of these factors in bacterial adherence, invasion, and host cell subversion remains to be determined.

## Data availability

This Whole Genome Shotgun project has been deposited at DDBJ/ENA/GenBank under the accession JAVKVK000000000. The version described in this paper is version JAVKVK010000000

## Acknowledgments

This research was supported by Coordenação de Aperfeiçoamento de Pessoal de Nível Superior (CAPES) (grant 99999.009868/ 2014-03), National Council for Scientific and Technological Development (CNPq) (grant 141586/2013-3), Fundação de Amparo à Pesquisa do Estado de São Paulo (FAPESP) (grants 2011/12664-5; 2023/05910-7), and the National Institutes of Health (NIH) (grant AI053067). Publication fee was supported by FAPESP (grant 2017/14821-7). The funders had no role in study design, data collection and interpretation, or the decision to submit the work for publication.

## Author contribution

Conceptualization: TATG and RTH; Data Curation: TATG; Formal Analysis: FTR, RTH and ACMS; Funding acquisition: TATG; Investigation: FTR, ACMS; Project Administration: FTR; Resources: VS and TATG; Supervision: RTH, VS, and TATG; Validation: FTR, RTH and ACMS; Visualization: FTR and ACMS; Writing – original draft: ACMS; Writing – review and editing: ACMS, RTH, TATG. All the authors read the manuscript’s final version and agreed with the manuscript submission.

## Reference

1. Ori EL, Takagi EH, Andrade TS, Miguel BT, Cergole-Novella MC, Guth BEC, Hernandes RT, Dias RCB, Pinheiro SRS, Camargo CH, Romero EC, Dos Santos LF. 2019. Diarrhoeagenic *Escherichia coli* and *Escherichia albertii* in Brazil: Pathotypes and serotypes over a 6-year period of surveillance. Epidemiol Infect 147.

2. Gomes TAT, Elias WP, Scaletsky ICA, Guth BEC, Rodrigues JF, Piazza RMF, Ferreira LCS, Martinez MB. 2016. Diarrheagenic *Escherichia coli*. Brazilian Journal of Microbiology 47:3–30.

3. Trabulsi LR, Keller R, Tardelli Gomes TA. 2002. Typical and atypical enteropathogenic *Escherichia coli*. Emerg Infect Dis 8:508–513.

4. Hernandes RT, Elias WP, Vieira MAM, Gomes TAT. 2009. An overview of atypical enteropathogenic *Escherichia coli*. FEMS Microbiol Lett 297:137–149.

5. Gaytán MO, Martínez-Santos VI, Soto E, González-Pedrajo B. 2016. Type Three Secretion System in Attaching and Effacing Pathogens. Front Cell Infect Microbiol 6.

6. Mcdaniel TK, Jarvis KG, Donnenberg MS, Kaper JB. 1995. A genetic locus of enterocyte effacement conserved among diverse enterobacterial pathogens. Proc Natl Acad Sci U S A 92:1664–1668.

7. Bando SY, Andrade FB, Guth BEC, Elias WP, Moreira-Filho CA, Pestana De Castro AF. 2009. Atypical enteropathogenic *Escherichia coli* genomic background allows the acquisition of non-EPEC virulence factors. FEMS Microbiol Lett 299:22–30.

8. Afset JE, Anderssen E, Bruant G, Harel J, Wieler L, Bergh K. 2008. Phylogenetic backgrounds and virulence profiles of atypical enteropathogenic *Escherichia coli* strains from a case-control study using multilocus sequence typing and DNA microarray analysis. J Clin Microbiol 46:2280–2290.

9. Hernandes RT, Hazen TH, Dos Santos LF, Richter TKS, Michalski JM, Rasko DA. 2020. Comparative genomic analysis provides insight into the phylogeny and virulence of atypical enteropathogenic *Escherichia coli* strains from Brazil. PLoS Negl Trop Dis 14:1–19.

10. Moreira FC, Vieira MAM, Ferreira AJP, Girão DM, Vaz TMI, Rosa ACP, Knobl T, Irino K, Freymüller E, Gomes TAT. 2008. *Escherichia coli* strains of serotype O51:H40 comprise typical and atypical enteropathogenic *E. coli* strains and are potentially diarrheagenic. J Clin Microbiol 46:1462–1465.

11. Gomes TAT, Irino K, Girão DM, Girão VBC, Guth BEC, Vaz TMI, Moreira FC, Chinarelli SH, Vieira MAM. 2004. Emerging enteropathogenic *Escherichia coli* strains? Emerg Infect Dis 10:1851–1855.

12. Arais LR, Barbosa A V., Andrade JRC, Gomes TAT, Asensi MD, Aires CAM, Cerqueira AMF. 2018. Zoonotic potential of atypical enteropathogenic *Escherichia coli* (aEPEC) isolated from puppies with diarrhoea in Brazil. Vet Microbiol 227:45– 51.

13. Vieira MAM, Andrade JRC, Trabulsi LR, Rosa ACP, Dias AMG, Ramos SRTS, Frankel G, Gomes TAT. 2001. Phenotypic and genotypic characteristics of *Escherichia coli* strains of non-enteropathogenic *E. coli* (EPEC) serogroups that carry EAE and lack the EPEC adherence factor and Shiga toxin DNA probe sequences. J Infect Dis 183:762–772.

14. Sampaio SCF, Gomes TAT, Pichon C, Du Merle L, Guadagnini S, Abe CM, Sampaio JLM, Le Bouguénec C. 2009. The flagella of an atypical enteropathogenic *Escherichia coli* strain are required for efficient interaction with and stimulation of interleukin-8 production by enterocytes in vitro. Infect Immun 77:4406–4413.

15. Sampaio S, Andrade J, Sampaio J, Carneiro C, Freymüller E, Gomes T. 2011. Distinct Interaction of Two Atypical Enteropathogenic *Escherichia coli* Strains with Enterocytes In Vitro. Open Microbiol J 5:65–71.

16. Liberatore AMA, Moreira FC, Gomes TAT, Menchaca-Diaz JL, Koh IHJ. 2011. Typical and atypical enteropathogenic *Escherichia coli* bacterial translocation associated with tissue hypoperfusion in rats. Brazilian Journal of Medical and Biological Research 44:1018–1024.

17. Sampaio SCF, Moreira FC, Liberatore AMA, Vieira MAM, Knobl T, Romão FT, Hernandes RT, Ferreira CSA, Ferreira AP, Felipe-Silva A, Sinigaglia-Coimbra R, Koh IHJ, Gomes TAT. 2014. Analysis of the Virulence of an Atypical Enteropathogenic *Escherichia coli* Strain *In Vitro* and *In Vivo* and the Influence of Type Three Secretion System. Biomed Res Int 2014:797508.

18. Riedl J, Crevenna AH, Kessenbrock K, Yu JH, Neukirchen D, Bista M, Bradke F, Jenne D, Holak TA, Werb Z, Sixt M, Wedlich-Soldner R. 2008. Lifeact: a versatile marker to visualize F-actin. Nat Methods 5:605.

19. Gruber CC, Sperandio V. 2014. Posttranscriptional Control of Microbe-Induced Rearrangement of Host Cell Actin. mBio 5.

20. Pacheco VCR, Yamamoto D, Abe CM, Hernandes RT, Mora A, Blanco J, Gomes TAT. 2014. Invasion of differentiated intestinal Caco-2 cells is a sporadic property among atypical enteropathogenic *Escherichia coli* strains carrying common intimin subtypes. Pathog Dis 70:167–175.

21. Rocha SPD, Abe CM, Sperandio V, Bando SY, Elias WP. 2011. Atypical enteropathogenic *Escherichia coli* that contains functional locus of enterocyte effacement genes can be attaching-and-effacing negative in cultured epithelial cells. Infect Immun 79:1833–1841.

22. Romão FT, Martins FH, Hernandes RT, Ooka T, Santos FF, Yamamoto D, Bonfim-Melo A, Jones N, Hayashi T, Elias WP, Sperandio V, Gomes TAT. 2020. Genomic Properties and Temporal Analysis of the Interaction of an Invasive *Escherichia albertii* With Epithelial Cells. Front Cell Infect Microbiol 10:749.

23. Müller D, Hagedorn P, Brast S, Heusipp G, Bielaszewska M, Friedrich AW, Karch H, Schmidt MA. 2006. Rapid identification and differentiation of clinical isolates of enteropathogenic *Escherichia coli* (EPEC), atypical EPEC, and Shiga toxin-producing *Escherichia coli* by a one-step multiplex PCR method. J Clin Microbiol 44:2626–2629.

24. Walters M, Sircili MP, Sperandio V. 2006. AI-3 synthesis is not dependent on *luxS* in *Escherichia coli*. J Bacteriol 188:5668–5681.

25. Koren S, Walenz BP, Berlin K, Miller JR, Bergman NH, Phillippy AM. 2017. Canu: scalable and accurate long-read assembly via adaptive k-mer weighting and repeat separation. Genome Res 27:722–736.

26. Vaser R, Sović I, Nagarajan N, Šikić M. 2017. Fast and accurate de novo genome assembly from long uncorrected reads. Genome Res 27:gr.214270.116.

27. Malberg Tetzschner AM, Johnson JR, Johnston BD, Lund O, Scheutz F. 2020. *In Silico* Genotyping of *Escherichia coli* Isolates for Extraintestinal Virulence Genes by Use of Whole-Genome Sequencing Data. J Clin Microbiol 58.

28. Joensen KG, Scheutz F, Lund O, Hasman H, Kaas RS, Nielsen EM, Aarestrup FM. 2014. Real-time whole-genome sequencing for routine typing, surveillance, and outbreak detection of verotoxigenic *Escherichia coli*. J Clin Microbiol 52:1501– 1510.

29. Camacho C, Coulouris G, Avagyan V, Ma N, Papadopoulos J, Bealer K, Madden TL. 2009. BLAST+: architecture and applications. BMC Bioinformatics 10.

30. Zankari E, Allesøe R, Joensen KG, Cavaco LM, Lund O, Aarestrup FM. 2017. PointFinder: a novel web tool for WGS-based detection of antimicrobial resistance associated with chromosomal point mutations in bacterial pathogens. Journal of Antimicrobial Chemotherapy 72:2764–2768.

31. Bortolaia V, Kaas RS, Ruppe E, Roberts MC, Schwarz S, Cattoir V, Philippon A, Allesoe RL, Rebelo AR, Florensa AF, Fagelhauer L, Chakraborty T, Neumann B, Werner G, Bender JK, Stingl K, Nguyen M, Coppens J, Xavier BB, Malhotra-Kumar S, Westh H, Pinholt M, Anjum MF, Duggett NA, Kempf I, Nykäsenoja S, Olkkola S, Wieczorek K, Amaro A, Clemente L, Mossong J, Losch S, Ragimbeau C, Lund O, Aarestrup FM. 2020. ResFinder 4.0 for predictions of phenotypes from genotypes. J Antimicrob Chemother 75:3491–3500.

32. Carattoli A, Zankari E, García-Fernández A, Voldby Larsen M, Lund O, Villa L, Møller Aarestrup F, Hasman H. 2014. In Silico Detection and Typing of Plasmids using PlasmidFinder and Plasmid Multilocus Sequence Typing. Antimicrob Agents Chemother 58:3895–3903.

33. Wirth T, Falush D, Lan R, Colles F, Mensa P, Wieler LH, Karch H, Reeves PR, Maiden MCJ, Ochman H, Achtman M. 2006. Sex and virulence in *Escherichia coli*: An evolutionary perspective. Mol Microbiol 60:1136–1151.

34. Liu B, Zheng D, Jin Q, Chen L, Yang J. 2018. VFDB 2019: a comparative pathogenomic platform with an interactive web interface. Nucleic Acids Res 47:687–692.

35. Zhou Z, Alikhan NF, Mohamed K, Fan Y, Achtman M. 2020. The EnteroBase user’s guide, with case studies on *Salmonella* transmissions, *Yersinia pestis* phylogeny, and *Escherichia* core genomic diversity. Genome Res 30:138–152.

36. Wattam AR, Davis JJ, Assaf R, Boisvert S, Brettin T, Bun C, Conrad N, Dietrich EM, Disz T, Gabbard JL, Gerdes S, Henry CS, Kenyon RW, Machi D, Mao C, Nordberg EK, Olsen GJ, Murphy-Olson DE, Olson R, Overbeek R, Parrello B, Pusch GD, Shukla M, Vonstein V, Warren A, Xia F, Yoo H, Stevens RL. 2017. Improvements to PATRIC, the all-bacterial Bioinformatics Database and Analysis Resource Center. Nucleic Acids Res 45:D535–D542.

37. Hazen TH, Donnenberg MS, Panchalingam S, Antonio M, Hossain A, Mandomando I, Ochieng JB, Ramamurthy T, Tamboura B, Qureshi S, Quadri F, Zaidi A, Kotloff KL, Levine MM, Barry EM, Kaper JB, Rasko DA, Nataro JP. 2016. Genomic diversity of EPEC associated with clinical presentations of differing severity. Nat Microbiol 1.

38. Letunic I, Bork P. 2021. Interactive Tree Of Life (iTOL) v5: an online tool for phylogenetic tree display and annotation. Nucleic Acids Res 49:W293–W296.

39. Gaytán MO, Martínez-Santos VI, Soto E, González-Pedrajo B. 2016. Type three secretion system in attaching and effacing pathogens. Front Cell Infect Microbiol 6:215590.

40. Pais SV, Kim E, Wagner S. 2023. Virulence-associated type III secretion systems in Gram-negative bacteria. Microbiology (United Kingdom) 169:001328.

41. Slater SL, Sågfors AM, Pollard DJ, Ruano-Gallego D, Frankel G. 2018. The type III secretion system of pathogenic escherichia coli. Curr Top Microbiol Immunol 416:51–72.

42. Ruano-Gallego D, Sanchez-Garrido J, Kozik Z, Núñez-Berrueco E, Cepeda-Molero M, Mullineaux-Sanders C, Clark JNB, Slater SL, Wagner N, Glegola-Madejska I, Roumeliotis TI, Pupko T, Fernández LÁ, Rodríguez-Patón A, Choudhary JS, Frankel G. 2021. Type III secretion system effectors form robust and flexible intracellular virulence networks. Science (1979) 371.

43. Campbell-Valois FX, Schnupf P, Nigro G, Sachse M, Sansonetti PJ, Parsot C. 2014. A fluorescent reporter reveals on/off regulation of the shigella type III secretion apparatus during entry and cell-to-cell spread. Cell Host Microbe 15:177–189.

44. Campbell-Valois FX, Pontier SM. 2016. Implications of Spatiotemporal Regulation of Shigella flexneri Type Three Secretion Activity on Effector Functions: Think Globally, Act Locally. Front Cell Infect Microbiol 6:28.

45. Van Nhieu GT, Guignot J. 2009. When *Shigella* Tells the Cell to Hang On. J Mol Cell Biol 1:64–65.

46. Leverton LQ, Kaper JB. 2005. Temporal Expression of Enteropathogenic *Escherichia coli* Virulence Genes in an *In Vitro* Model of Infection. Infect Immun 73:1034–1043.

47. Shimizu T, Ichimura K, Noda M. 2016. The surface sensor NlpE of enterohemorrhagic *Escherichia coli* contributes to regulation of the type III secretion system and flagella by the Cpx response to adhesion. Infect Immun 84:537–549.

